# Impact of Neurons on Group B *Streptococcus* Interactions at the Blood Brain Barrier

**DOI:** 10.64898/2026.07.31.741976

**Authors:** Lena M. Seyfarth, Natalie G. Alexander, Alyssa S. Arnett, Kathryn N. Seely, Benjamin T. Klemp, Taryn E. Keyzer, Nadine Vollmuth, Caylah Griffin, Michael D. Burton, Erica L. Sanchez, Brandon J. Kim

**Author notes:** Address correspondence to Brandon J. Kim. Lena M. Seyfarth and Natalie G. Alexander contributed equally to this work. Author order was determined alphabetically by first name.

## Abstract

*Streptococcus agalactiae* (Group B *Streptococcus*, GBS) is a Gram-positive opportunistic pathogen and the leading cause of neonatal bacterial meningitis, a life-threatening infection of the central nervous system (CNS) that occurs when bacteria cross the blood-brain barrier (BBB). GBS commonly colonizes the maternal genital tract and is a major cause of invasive neonatal disease, including bacteremia and meningitis. Despite treatment advances, GBS meningitis remains associated with substantial mortality and long-term neurological sequelae. The BBB is a highly specialized barrier formed by brain endothelial cells (BECs) that restrict microbial entry into the CNS through tightly regulated intercellular junctions. The BBB exists within the neurovascular unit, where neurons and other CNS cell types actively regulate endothelial barrier properties through intercellular signaling. However, the contribution of neuronal-endothelial interactions to BBB function during neonatal meningitis remains poorly understood. To investigate the mechanisms by which GBS disrupts and penetrates the BBB, we utilized induced pluripotent stem cell (iPSC)-derived brain-like endothelial cells. EZ-Sphere-derived neurons generated from the same iPSC source were incorporated into an isogenic BBB model to determine whether neuronal-endothelial communication influences GBS interaction with BECs. Neuronal co-culture significantly reduced GBS adherence to and invasion of BECs while preserving tight junction integrity during infection. Application of neuron-conditioned medium similarly decreased bacterial adherence and invasion, suggesting that neuron-derived soluble factors enhance barrier integrity during GBS infection. Together, these findings demonstrate that neuronal signaling enhances BBB resistance to GBS and highlight a previously underappreciated role for neurovascular crosstalk in limiting bacterial pathogenesis.

**Importance:** Group B *Streptococcus* (GBS) is the leading cause of bacterial meningitis in newborns. GBS interacts with and crosses the blood-brain barrier (BBB) contributing to a potentially fatal infection without treatment. Understanding how the BBB interacts with bacterial pathogens is critical for developing new strategies to protect vulnerable infants. In this study, we used human stem cell-derived models to recapitulate the BBB and examine how communication between brain endothelial cells and neurons influence host response to bacterial infection. These findings identify neuronal-endothelial communication as a potential contributor to BBB protection and provide a foundation for future studies aimed at preventing GBS invasion of the central nervous system.

## Introduction

Bacterial meningitis is a life-threatening infection of the central nervous system (CNS) that occurs when pathogens penetrate the blood-brain barrier (BBB) (1–4). Disease burden is greatest among young children, particularly neonates (5). Despite antimicrobial therapy, neonatal meningitis remains associated with substantial mortality, and up to 50% of surviving neonates develop permanent neurological sequelae (3, 5, 6). Group B *Streptococcus* (GBS, *Streptococcus agalactiae)* is the leading cause of neonatal bacterial meningitis worldwide. GBS asymptomatically colonizes the female genital tract and remains a major cause of neonatal invasive disease, including bacteremia and meningitis (7). Successful CNS invasion requires survival in the bloodstream followed by traversal of the BBB, although the host factors that influence BBB susceptibility to GBS infection remain incompletely understood.

The BBB is a highly specialized cellular barrier that makes up the cerebral vasculature formed by brain endothelial cells (BECs), which restrict microbial entry through tightly regulated intercellular junctions and selective transport mechanisms (8–10). BECs function within the neurovascular unit (NVU), a multicellular structure composed of endothelial cells, pericytes, astrocytes, microglia, and neurons that collectively regulate BBB function and CNS homeostasis (11–13). The barrier properties of the BBB are maintained by tight junction proteins including Claudins and Occludin, which are anchored by zonula occludens (ZO) scaffold proteins and restrict paracellular permeability (11, 14). Genetic disruption of critical tight junction proteins, such as claudin-5, results in severe barrier dysfunction and perinatal lethality in mice, highlighting the importance of junctional integrity for CNS protection (15, 16).

*In vitro* studies of the BBB have previously relied on primary and immortalized endothelial cell models; however, both systems have notable limitations (17, 18). Primary cells rapidly lose key BBB characteristics upon removal from the brain microenvironment, whereas immortalized lines exhibit reduced barrier integrity, including low transendothelial electrical resistance (TEER) and disorganized tight junctions (17, 19, 20). Advances in induced pluripotent stem cell (iPSC) technology have enabled the development of renewable BBB models that more closely recapitulate *in vivo* barrier properties (21–24). Importantly, iPSC-derived systems permit generation of multiple CNS cell types from a common genetic background, enabling development of isogenic NVU models that more closely replicate cellular interactions within the BBB (23, 25). While the contributions of endothelial cells, astrocytes, and pericytes to BBB maintenance have been well characterized, little is known about whether these cells influence BBB susceptibility to bacterial infection.

Previous studies investigating neonatal meningitis have shown that GBS employs multiple adhesins, invasins, and secreted toxins to promote interaction with brain endothelial cells and disrupt BBB integrity (26–29). These investigations have largely focused on direct pathogen-endothelial interactions and have identified several bacterial and host pathways that contribute to CNS invasion (30–32). However, these approaches do not account for the broader cellular environment of the NVU, and it remains unclear whether neighboring cells influence endothelial susceptibility to GBS infection. Because neurons actively contribute to BBB development and maintenance, neuron-endothelial communication may represent an unrecognized determinant of BBB resistance during bacterial meningitis. Here we explore the contribution of neurons to GBS-BEC interactions and begin investigating the role of the NVU to bacterial-BBB disruption.

In this study, we combined established iPSC differentiation protocols to generate an isogenic human BBB model comprising brain endothelial cells and neurons derived from the same donor line (23–25). Using this system, we investigated whether neuronal-endothelial communication influences GBS interactions with the BBB. We demonstrate that neuronal co-culture reduces GBS adherence to and invasion of brain endothelial cells while preserving barrier integrity during infection. Furthermore, neuron-conditioned medium recapitulates these protective effects, suggesting that neuron-derived soluble factors enhance BBB protection to GBS. Together, these findings identify neuronal-endothelial crosstalk as a previously unrecognized determinant of host response at the BBB and establish a foundation for defining neurovascular mechanisms that limit bacterial CNS invasion.

## Materials and Methods

### Bacterial Strains and Preparation

The hypervirulent wild-type GBS clinical isolate COH1 (serotype III, multilocus sequence type 17 [MLST-17]) was used in all experiments (1). GBS was grown overnight in Todd-Hewitt broth (THB) at 37°C, subcultured the following day, and grown to mid-logarithmic phase (OD_600_ = 0.4-0.6). Bacterial cultures were pelleted by centrifugation and resuspended in 250 uL phosphate-buffered saline (PBS). The bacterial suspension was adjusted to an OD_600_ of 0.4 and subsequently diluted 1:10 in appropriate endothelial cell medium prior to infection. All infection assays were performed at a multiplicity of infection (MOI) of 10.

### Cell Lines

IMR90-4 iPSCs (WiCell) were maintained on Matrigel-coated (Corning 354234) 6-well tissue culture plates in StemFlex medium (Gibco A3349401), with medium changes performed daily. iPSCs were passaged twice per week once at 80% confluence.

Immortalized hCMEC/D3 BECs were maintained in EndoGro MV medium (Millipore SCME004) on plates coated with 1% rat-tail collagen (RTC, ThermoFisher A1048301) as previously described (33, 34). SH-SY5Y cells were maintained in 89% DMEM without phenol red (ThermoFisher 21063-029), 10% Fetal Bovine Serum (FBS), and 1% penicillin-streptomycin (Gibco 15140122). SH-SY5Y cells were passaged once at 80% confluence, with medium changes performed every other day.

### Human Serum iPSC-derived Brain-like Endothelial Cell (iBEC) differentiation

IMR90-4 iPSCs were differentiated into iBECs as previously described (24, 35). Briefly, a single-cell suspension of iPSCs was seeded onto Matrigel-coated cell culture flasks at a density of 10,000 cells/cm^2^ and grown for two days. Differentiation was initiated by replacing StemFlex medium with unconditioned medium (UM) consisting of DMEM/F12 (ThermoFisher 11320033), 20% KnockOut Serum Replacement (KOSR; Gibco 10828028), 1% nonessential amino acids (NEAA; Invitrogen 11140-050), 0.5% GlutaMAX (Invitrogen 35050-061), and 0.0007% β-Mercaptoethanol (ß-ME; Sigma M3148). Cells were maintained in UM for 6 days with daily medium changes. After this, the media was transitioned to EC +/+ medium consisting of Human Endothelial Serum Free Medium (hESFM; Gibco 11111044), 1% human serum from platelet-poor derived plasma (Sigma P2918), and 20 ng/mL basic fibroblast growth factor (bFGF; Peprotech 100-18B) for two days. iBECs were subsequently purified onto fibronectin (Sigma F1141) and collagen IV (Sigma C5533) coated plates or Transwell inserts using EC -/- medium.

### EZ-Sphere and iNeuron Differentiation

EZ-Spheres were generated from IMR90-4 iPSCs as previously described (23, 36). Briefly, iPSCs were dissociated with Versene (ThermoFisher 15040066) for 7 min. Cells were collected by centrifugation and resuspended in EZ-Sphere maintenance medium consisting of high-glucose DMEM (ThermoFisher 11965092), F12 (ThermoFisher 11765062), 1% antibiotic-antimycotic (ThermoFisher 15240062), 2% B-27 supplement minus vitamin A (ThermoFisher 12587010), 2 μg/mL heparin (Sigma H3149), 100 ng/mL epidermal growth factor (EGF, Millipore GF144), and 100 ng/mL bFGF. Cells were maintained in ultra-low attachment flasks to promote sphere formation, and free-floating EZ-Spheres formed within 1-2 weeks of culture. To maintain sphere size and viability, EZ-Spheres were mechanically passaged weekly using a McIlwain Tissue Chopper (Model TC752).

EZ-Spheres were subsequently differentiated into induced neurons (iNeurons) according to previously established protocols (23). Briefly, EZ-Spheres were dissociated into single cells by incubation with Accutase (Innovative Cell Technologies AT104-500) for 10-30 min, followed by gentle trituration to generate a single-cell suspension. Enzymatic dissociation was quenched with four volumes of DMEM/F12, and viable cells were quantified using trypan blue exclusion (Gibco 15250061). Cells were subsequently pelleted by centrifugation at 1000 rpm for 10 min, and the supernatant was aspirated.

The cell pellet was resuspended in EZ-Sphere maintenance medium at a density of 1 million cells/mL. Cells were seeded onto Matrigel-coated culture plates at a density of 25,000 cells/cm^2^. After 24 h, the medium was replaced with neuronal differentiation medium consisting of DMEM/F12 supplemented with 1% penicillin-streptomycin, 2% B-27 supplement minus vitamin A, and 2 ug/mL heparin. Medium was changed every 48 h, and iNeurons were differentiated for 14 days prior to experimental use. Neuronal differentiation was confirmed by βIII-tubulin expression.

### Initiation of BEC-Neuron Co-Culture Experiments

One day post purification, iBECs were maintained as either monocultures or in indirect co-culture with iNeurons in EC-/- medium (23). To enhance iBEC attachment and retention, 10 μM ROCK inhibitor Y-27632 (Tocris 1254) was added to the culture medium during the first 24 h of co-culture. hCMEC/D3 cells were cultured in EndoGro medium either as monocultures or in indirect co-culture with iNeurons beginning 4 days post seeding. All experiments were performed 24 h after establishment of co-culture conditions.

### Initiation of Neuron Conditioned Medium BEC Experiments

Neuron-conditioned medium was generated by culturing iNeurons in corresponding endothelial cell medium for 48 h. Following conditioning, the medium was clarified by centrifugation and passed through a 0.22μm filter and stored at -20°C until use. For conditioned medium experiments, BECs were pretreated for 24 h with either control endothelial cell medium or a 1:1 mixture of neuron-conditioned medium and control medium prior to GBS infection. Two hours before infection, cultures were switched to either fresh control medium or 100% neuron-conditioned medium, respectively, and maintained in these conditioned throughout the infection period.

### TEER measurement

iBECs were purified onto a collagen IV- and fibronectin-coated 12-well Transwell inserts as previously described(22, 24). TEER was measured 24 h after iBEC purification using an EVOM Manual Epithelial Voltohmeter (World Precision Instruments, EVM-MT-03-02). For hCMEC/D3s, TEER was measured on the day of experimental use. For conditioned medium experiments, TEER measurements were obtained 24 h after treatment with neuron-conditioned medium or control medium.

### Isolation of Primary Dorsal Root Ganglia (DRG) Tissue

Primary peripheral DRG neurons were isolated from thoracic and lumbar spinal levels (T10-T13, L1-L6) of male and female C57BL/6J mice, (n=5; Jackson Laboratory, stock number 000664) that were bred in-house. Mice were housed at a maximum of 5 animals per cage and given *ad libitum* access to food and water. Animals were housed in a temperature-controlled facility kept at 21°-23°C and 50% humidity and maintained on a 12 h light cycle. Animals were 6-10 weeks of age at the time of tissue collection. Mice were deeply anesthetized with isoflurane and euthanized by decapitation. DRGs were collected and immediately placed in chilled Hank’s Balanced Salt Solution (HBSS; Cytiva, SH30588.01). DRGs were enzymatically digested by incubation in collagenase A (1:1;A (Sigma-Aldrich, 10103586001) for 20 min at 37°C and collagenase D (1:1:10%; D (Sigma-Aldrich, 1188866001): HBSS: papain (Sigma-Aldrich, 10108014001)) for 20 min at 37°C and Trypsin Inhibitor solution (1:1:1; (Sigma-Aldrich, 10109886001): HBSS:Media) to stop the digestion reaction. Cells were triturated and filtered with a 70µm cell strainer (Corning, 431751), centrifuged, and resuspended in 300 µl of DMEM/F12 medium (Cytiva, SH30023.FS) supplemented with 2% Penicillin Streptomycin (ThermoFisher, 15070063) and 10% FBS (Cytiva, SH30088.03IR25-40). Neurons were counted with TC 20 Automated Cell

Counter (BioRad) with trypan blue exclusion. Glass coverslips (12mm, #1; Fisher Scientific, 08-774-384) were coated with 2 µg/mL Poly-D-Lysine solution (Sigma-Aldrich, P0899) and placed into 12-well culture plates (Corning, 07-200-82). A total of 30,000 cells were seeded onto each coverslip in a 100 µl droplet on the center of the coverslip and incubated at 37°C for 2 h to allow for cell attachment. Following 2 h incubation, wells were filled with DMEM/F12 media supplemented with 10% FBS, 2% Penicillin Streptomycin, and 10 ng/mL Nerve Growth Factor (Thomas Scientific, C970E23).

### GBS Adherence and Invasion Assay

iBECs were purified onto collagen IV- and fibronectin-coated 12-well Transwell inserts for co-culture experiments, while hCMEC/D3 cells were seeded onto 1% RTC-coated 12-well Transwell inserts. Bacterial adherence and invasion assays were performed as previously described (33). For adherence assays, BECs were infected with WT COH1 at a MOI of 10 for 30 min at 37°C + 5% CO_2_. Cells were washed five times with PBS, detached with 0.25% Trypsin-EDTA, and lysed in 0.025% Triton-X 100 in PBS. Serial dilutions of the lysates were plated on THB agar to quantify adherent bacteria. For Invasion assays, BECs were infected with WT COH1 at a MOI of 10 for 2 h at 37°C + 5% CO_2_ and subsequently washed once with PBS. Extracellular bacteria were eliminated by incubation in EC -/- medium containing gentamicin (100 μg/mL) for 2 h at 37°C in 5% CO₂. BECs were detached with Trypsin-EDTA and lysed in 0.025% Triton X- 100 in PBS. Serial dilutions of the lysates were plated on THB agar to quantify intracellular bacteria.

### qRT-PCR

iBECs and hCMEC/D3 cells were infected with WT COH1 or maintained as uninfected controls for 4 h at 37°C + 5% CO_2_. For iNeuron co-culture experiments, Transwell inserts were transferred to a new 12-well plate following bacterial infection to isolate the BECs. Total RNA was isolated using the NucleoSpin RNA kit (Macherey-Nagel MANA740955.25). cDNA was synthesized using the qScript cDNA Synthesis Kit (Quantabio 95047-500) according to the manufacturer’s instructions. Unless otherwise indicated, 300 ng of total RNA was used as input for each cDNA synthesis reaction.

Quantitative real-time PCR (qPCR) was performed using SYBR Green Master Mix (ThermoFisher, A25743) to assess the expression of *GAPDH* (glyceraldehyde-3-phosphate dehydrogenase), *OCLN* (Occludin), *TJP1* (tight junction protein 1; ZO-1), *CLDN5* (claudin-5), *SNAI1* (Snail1), and *TUBB3* (βIII-tubulin). Primer sequences were obtained from previously published studies (1, 37). Amplification and data acquisition were performed using a QuantStudio 3 Real-Time PCR System (ThermoFisher). Relative gene expression was calculated using the comparative cycle threshold (ΔΔC_T_) method, with GAPDH as the endogenous reference gene. Data are presented as fold change relative to the uninfected control condition.

### Western Blot

Protein concentrations were quantified using a BCA Protein Assay kit (ThermoFisher 23225) according to the manufacturer’s instructions. Equal concentrations of protein were loaded onto 4%-12% Tris-glycine SDS-PAGE gels and for 1 h at 110 V. Proteins were subsequently transferred to nitrocellulose membranes for 90 min at 300 mA. Following transfer, membranes were stained with Ponceau S to verify equal protein transfer and imaged using an Azure BioSystems multichannel imager. Membranes were then blocked in 5% (w/v) nonfat dry milk in TBST (Tris-buffered saline containing 0.1% Tween-20) for 1 h at room temperature. Membranes were incubated in primary antibodies overnight at 4°C (Table 1). The following day, membranes were washed three times with TBST for 5 min per wash before incubation with the appropriate secondary antibodies for 1 h at room temperature. Immunoreactive bands were detected using enhanced chemiluminescence substrate and imaged using an Azure Biosystems chemiluminescent imaging system.

**Table 1:** Antibodies used in this study.

| Antibody | Species | Source and Product Number |
| --- | --- | --- |
| IgG1 Anti-Occludin | Mouse | ThermoFisher 33-1500 (OC-3F10) |
| IgG1 Anti-ZO1 | Mouse | ThermoFisher 33-9100 (ZO1-1A12) |
| IgG1 Anti-Claudin-5 | Mouse | ThermoFisher 35-2500 (4C3C2) |
| Anti-β-Tubulin III Antibody | Rabbit | Sigma T3952 |
| IgG1 Anti-GAPDH | Mouse | ThermoFisher AM3400 |

### Immunostaining and JAnaP

iBECs were infected with GBS or left uninfected as controls for 4 h at 37°C + 5% CO_2_. For monoculture and neuron co-culture experiments, Transwell inserts were transferred to new 12-well plates following the incubation period isolate iBECs. Following infection, iBECs were fixed and immunostained as previously described (33, 35). Briefly, iBECs were fixed in ice-cold 100% methanol and incubated with primary antibodies overnight at 4°C (Table 1). Primary isolated DRG neurons were stained with βIII-tubulin for confirmation of pure neuronal population. All samples were subsequently incubated with secondary antibodies (1:200 dilution) for a minimum of 1 h at room temperature. Images were acquired using a Nikon Eclipse Ti2 inverted microscope and processed using ImageJ software. Scale bars represent 100 μm. Tight Junction continuity was quantified using the Junction Analyzer Program (JAnaP) as previously described (38–40). Data are presented as the percentage of continuous junctions for ZO-1 and Occludin across experimental conditions.

### Statistical Methods

Statistical analysis and figure generation were performed using GraphPad Prism version 10.4.1. For pairwise comparisons, Student’s *t*-tests were performed where appropriate. For comparisons involving more than two groups, analysis of variance (ANOVA) was used unless otherwise specified. Data are presented as mean ± standard deviation (SD). Statistical significance was defined as *p* < 0.05.

## Results

### Neuronal co-culture reduces GBS Adherence to and Invasion of BECs

Previous studies have demonstrated that GBS adheres to and invades brain endothelial cells (1, 33, 41–44). To determine whether neuronal-endothelial interactions influence these processes, we compared GBS infection of iBECs cultured alone or indirectly cultured with iNeurons. Human serum-derived iBECs exhibited characteristic endothelial and barrier-associated markers, including VE-Cadherin, Occludin, ZO-1, and Claudin-5 consistent with previous studies (Supplement Figure 1) (23, 24, 45). EZ-Sphere-derived iNeurons (Figure 1A) expressed the neuronal marker βIII-tubulin, confirming successful neuronal identity and validating the co-culture model as previously described (Figure 1B-C) (23, 25). To determine whether neuronal co-culture influences endothelial barrier function, iBECs and hCMEC/D3 cells were indirectly co-cultured with iNeurons and TEER was measured. Co-culture with iNeurons significantly increased TEER in both endothelial models, indicating enhanced barrier integrity (Figure 1E-F). We next examined whether these changes in barrier integrity were associated with altered bacterial interactions. Confluent iBEC monolayers cultured alone or in indirect co-culture with iNeurons were infected with GBS. Compared with iBEC monocultures, iBEC-neuron co-cultures exhibited significantly lower levels of adherent and intracellular bacteria (Figure 1G-H), suggesting that a neuronal protective effect limit GBS association with and entry into endothelial cells. To determine whether this phenotype extended beyond iPSC-derived endothelial cells, we performed parallel experiments using the immortalized human brain endothelial cell line hCMEC/D3. Consistent with the findings in iBECs, neuronal co-culture significantly reduced GBS adherence and invasion of hCMEC/D3 cells relative to monoculture controls (Figure 1I-J). Together, these findings demonstrate that neuronal co-culture limits GBS adherence to and invasion of brain endothelial cells across multiple endothelial models, supporting a previously unrecognized role for neuron-endothelial crosstalk in regulating susceptibility to bacterial infection.

**Figure 1:**
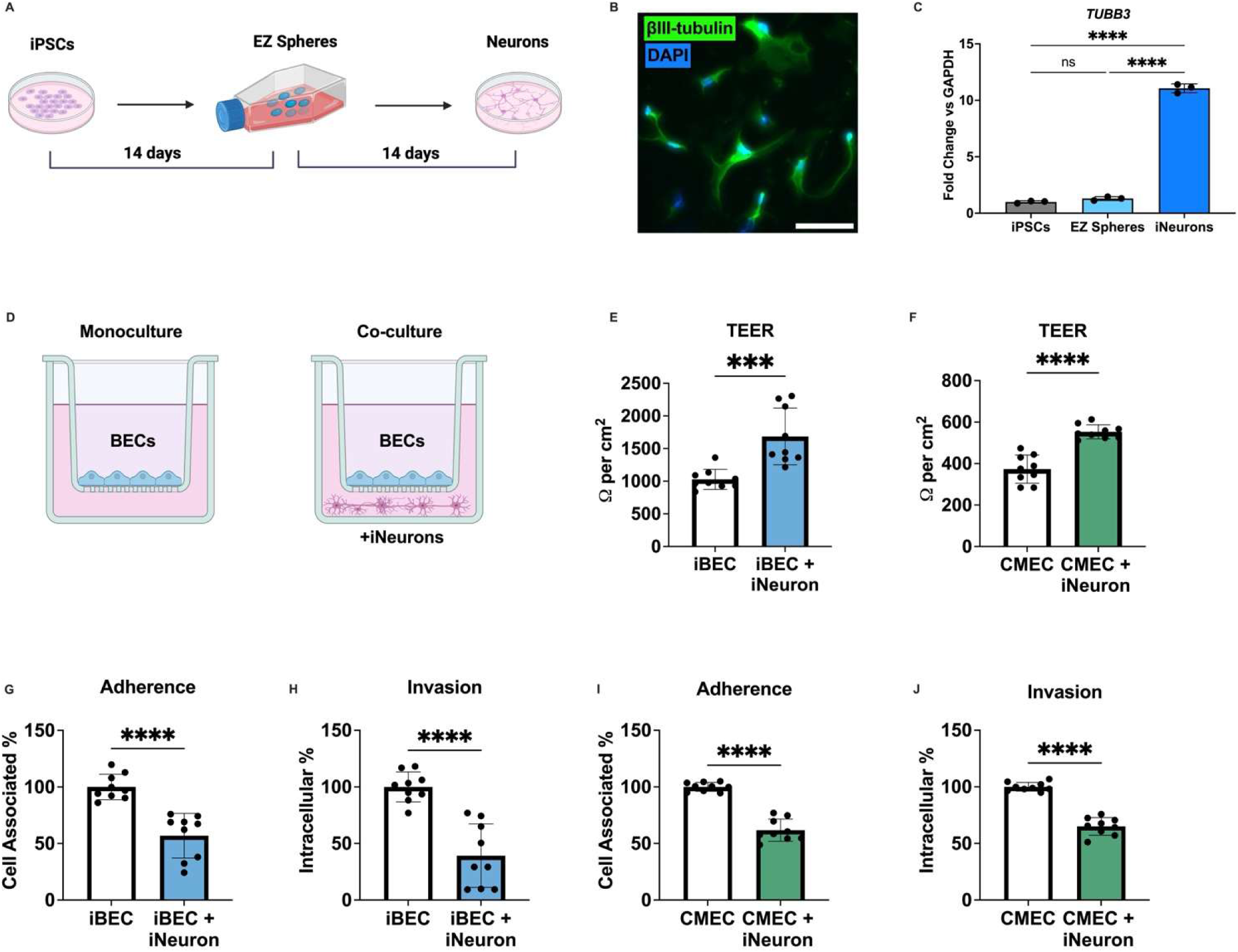
Neuronal co-culture preserves BBB integrity and reduces GBS adherence to and invasion of brain endothelial cells. (A) Schematic of differentiation of IMR90-4 induced pluripotent stem cells (iPSCs) into intermediate EZ-Spheres and subsequent generation of βIII-tubulin-positive induced neurons (iNeurons). (B) Representative immunofluorescence image showing βIII-tubulin expression in differentiated iNeurons. (C) qPCR of iPSCs, EZ Spheres, and iNeurons expression of *TUBB3*, *n* =3. (D) Schematic of BEC monoculture and BEC-iNeuron co-culture conditions. (E) TEER of iBECs and of (F) hCMEC/D3s showing neurons improve TEER similar to previous observations. (G) Adherence of wild-type (WT) Group B Streptococcus (GBS) strain COH1 to iBECs cultured alone or in co-culture with iNeurons. (H) Invasion of WT GBS COH1 into iBECs cultured alone or in co-culture with iNeurons. (I) Adherence and (J) invasion of WT COH1 to hCMEC/D3 cells following infection at MOI of 10. Adherence and invasion normalized to infected monoculture conditions. . Statistical significance for adherence and invasion assays was determined using Student’s *t*-test. **** p<0.0001, ns, non-significant. Scale bar is 100 uM. iBEC experiments were performed across three independent differentiations in technical triplicate, *n* = 9. hCMEC/D3 experiments were performed across three independent passages in technical triplicate, *n* = 9. Normality tests were performed and data are presented as mean ± SD

#### Neuronal Secreted Factors Impact Bacterial Adherence to and Invasion of iBECs

Given that neuronal co-culture reduced GBS adherence to and invasion of brain endothelial cells, we aimed to see whether this protective effect required live neuron-endothelial contact or could be mediated by neuron-derived soluble factors. To address this question, conditioned medium generated from EZ-Sphere-derived neurons was applied to endothelial cultures prior to GBS infection (Figure 2A). Neuron-conditioned medium was generated by incubating differentiated iNeurons in endothelial cell culture medium for 48 h before collection and sterile filtration (Figure 2A). To evaluate its functional effects, iBECs were pretreated with either control medium or neuron-conditioned medium prior to infection. Exposure to neuron-conditioned medium was maintained throughout the infection period. Following infection with GBS, pretreatment with neuron-conditioned medium resulted in a significant reduction in bacterial adherence to iBECs compared to cells maintained in control medium (Figure 2B).

**Figure 2:**
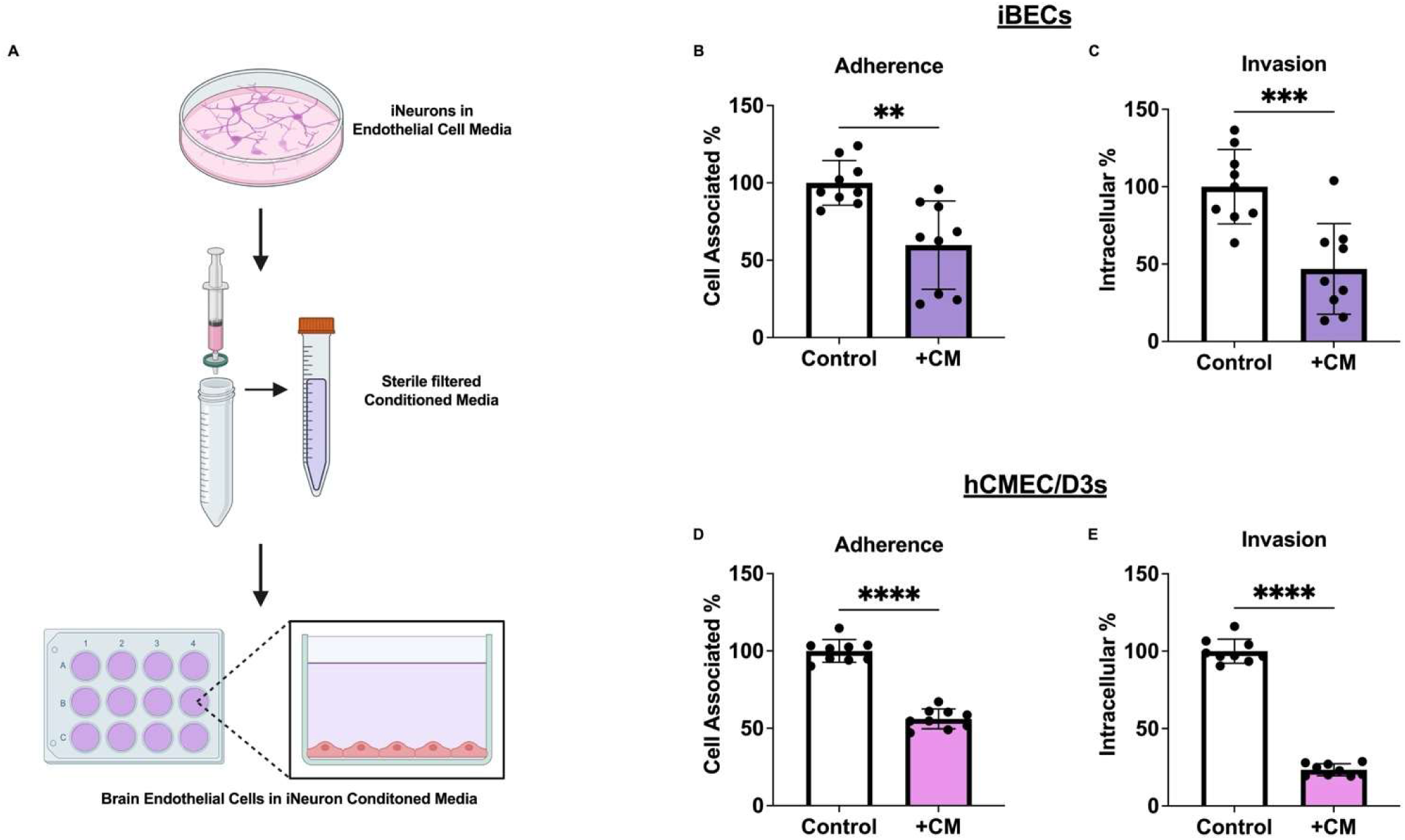
Neuron-conditioned medium reduces GBS adherence to and invasion of BECs. (A) Experimental schematic of neuron-conditioned medium generation and treatment. EZ-Sphere-derived neurons were cultured in endothelial cell medium for 48 h prior to conditioned medium collection and application to endothelial cells. (B-C) Adherence and invasion of wild-type (WT) Group B *Streptococcus* (GBS) strain COH1 in iBECs following treatment with control or neuron-conditioned medium. (D-E) Adherence and invasion of WT GBS COH1 in hCMEC/D3 cells following treatment with control or neuron-conditioned medium. Bacterial recovery was normalized to the corresponding control medium condition. Treatment with neuron-conditioned medium reduced both bacterial adherence and intracellular invasion in iBECs and hCMEC/D3 cells, indicating that neuron-derived soluble factors are sufficient to recapitulate the protective effects observed in neuronal co-culture. These findings indicate that direct neuron-endothelial contact is not required for protection against GBS and suggest that soluble neuron-derived signals contribute to BBB protection. Data are presented as mean ± SD from three independent iBEC differentiations or hCMEC/D3 passages performed in technical triplicate (*n* = 9). Statistical significance was determined using Student’s *t*-test. **p < 0.01, ***p < 0.001, ****p < 0.0001.

Similarly, intracellular invasion was markedly decreased in iBECs exposed to conditioned medium (Figure 2C), demonstrating that neuron-derived soluble factors are sufficient to recapitulate the protective phenotype observed in neuronal co-culture. To determine whether this effect was extended across endothelial models, we performed parallel experiments with hCMEC/D3s. Consistent with the iBEC findings, neuron-conditioned EndoGro medium significantly reduced both GBS adherence to and invasion of hCMEC/D3 cells (Figure 2D-E). We additionally assessed endothelial barrier function by measuring TEER following treatment with conditioned medium generated from primary mouse dorsal root ganglion (DRG) neurons or SH-SY5Y neuroblastoma cells (Supplement Figure 2). Whereas SH-SY5Y conditioned medium significantly increased TEER, DRG neuron-conditioned medium decreased TEER, indicating that soluble factors released by distinct neuronal populations differentially regulate endothelial barrier properties. Notably, SH-SY5Y conditioned medium was sufficient to decrease bacterial adherence to iBECs (Supplement Figure 2). Together, these findings demonstrate that neuron-derived soluble factors are sufficient to limit GBS interaction with brain endothelial cells and suggest that soluble neuron-endothelial signaling contributes to host defense at the BBB. These data indicate that neurons actively modulate endothelial susceptibility to bacterial infection through secreted factors, identifying a previously unrecognized mechanism of neurovascular protection against GBS.

#### Neuronal Co-Culture preserves tight junction integrity during GBS infection

Previous studies have shown that GBS disrupts tight junction components in iBECs and impairs BBB barrier function (1, 37). We next investigated whether co-culture with iNeurons could mitigate GBS-induced tight junction disruption. Immunostaining of mock and GBS-infected iBECs in monoculture or neuron co-culture revealed preservation of continuous tight junction staining patterns in GBS-infected iBECs co-cultured with neurons (Figure 3A-B). To quantify these effects, we used the Junction Analyzer Program (JAnaP) to assess the continuity of tight junction proteins ZO-1 and Occludin, as previously described (38–40, 46). Quantitative image analysis using JAnaP demonstrated significantly reduced junctional fragmentation and increased continuity of both ZO-1 (Figure 3C) and Occludin (Figure 3D) in GBS-infected iBEC-neuron co-cultures compared with infected iBEC monocultures. Expression of *SNAI1*, which encodes the transcriptional repressor Snail1 and has previously been shown to be associated with tight junction loss and was increased in GBS-infected iBEC monocultures but not in infected iBEC-neuron co-cultures (Figure 3E) (37). To assess the impact of neuronal co-culture on BBB function during infection, TEER was measured following GBS infection. GBS infection reduced TEER in iBEC monocultures, indicating disruption of barrier integrity. In contrast, iBECs maintained in co-culture with neurons exhibited significantly higher TEER values following infection, demonstrating preservation of barrier function (Supplement Figure 3A). Similar protective effects were observed in hCMEC/D3 co-cultures, where neuronal presence attenuated the infection-associated decline in TEER compared with infected monocultures (Supplement Figure 3B). Together, these data indicate that neuronal co-culture preserves BBB integrity during GBS infection, potentially through suppression of *SNAI1*-associated pathways involved in tight junction repression.

**Figure 3:**
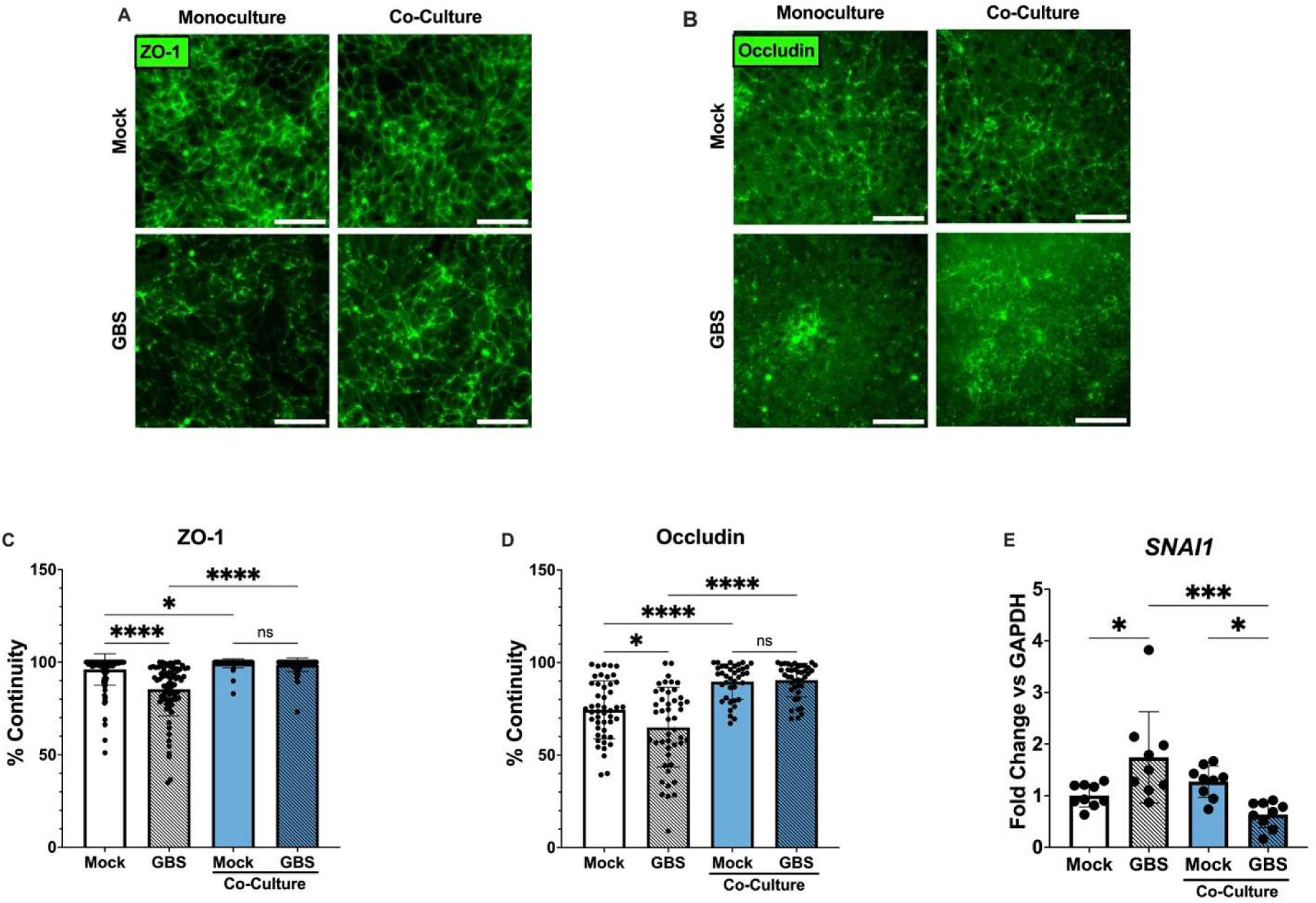
Neuronal Co-Culture preserves tight junction organization and barrier integrity during GBS infection. (A) Representative immunofluorescence images of iBECs maintained in monoculture or co-culture with iNeurons under mock and GBS-infected conditions. Cells were stained for the tight junction proteins ZO-1 (A) and Occludin (B) to assess junctional organization following infection. (C-D) Quantification of Occludin and ZO-1 junctional continuity using the Junction Analyzer Program (JAnaP). Neuronal co-culture preserved tight junction continuity in GBS-infected iBECs compared with infected monocultures. (E) Expression of SNAI1, a transcriptional repressor associated with tight junction disruption and barrier dysfunction. Neuronal co-culture prevented the infection-associated increase in *SNAI1* expression observed in iBEC monocultures. Together, these findings demonstrate that neuronal co-culture mitigates GBS-induced disruption of endothelial tight junctions and preserves BBB barrier integrity during infection. (C-D) Data are presented as percent continuity of individual cells from images acquired across three independent biological replicates in technical triplicate (C) *n* =69-118, (D) *n* = 38-47. (E) Data are presented as mean ± SD from three independent biological replicates in technical triplicate, *n* = 9. Ordinary one-way ANOVA was used to determine significance; *p < 0.05, ****p < 0.0001, ns, non-significant.

#### Neuron-conditioned medium preserves tight junction integrity during GBS infection

Given that neuronal co-culture preserved barrier integrity and reduced GBS-induced tight junction disruption, we next sought to determine whether these protective effects required live neuron-endothelial culture or could instead be mediated by neuron-derived soluble factors. To address this question, iBECs were treated with neuron-conditioned medium prior to mock or GBS infection. Immunostaining for the tight junction proteins ZO-1 and Occludin revealed that neuron-conditioned medium preserved continuous junctional staining patterns in GBS-infected iBECs relative to control-treated cells similar to our observations during co-culture (Figure 4A,C). Quantitative image analysis using JAnaP demonstrated significantly greater continuity of ZO-1 (Figure 4B) and Occludin (Figure 4D) staining in neuron-conditioned medium treated iBECs compared with controls. Because preservation of tight junction integrity by neuron-derived soluble factors during GBS infection has not previously been described, we next sought independent molecular validation of these findings. Consistent with the imaging data, immunoblot analysis demonstrated significantly increased ZO-1 and Occludin protein abundance in GBS-infected iBECs treated with neuron-conditioned medium compared with control-treated cells (Figure 4E-G). In contrast, neuron-conditioned medium treatment did not significantly alter ZO-1 and Occludin protein abundance in GBS-infected hCMEC/D3 cells (Supplement Figure 4), suggesting potential model-specific differences in endothelial responses to neuron-derived factors. To further validate these observations at the transcriptional level, we examined expression of barrier-associated tight junction genes including *OCLN* (Occludin) and *TJP1* (ZO-1) (Figure 4H-I). In addition, we assessed expression of *SNAI1*, a transcriptional repressor of tight junction expression. Notably, *SNAI1* expression was significantly upregulated in GBS-infected iBECs maintained in control medium but not in neuron-conditioned medium treated iBECs (Figure 4J). Together, these findings demonstrate that neuron-derived soluble factors are sufficient to preserve tight junction organization and promote barrier-protective responses in iBECs during GBS infection. These data support a model in which soluble neuron-endothelial signaling enhances BBB response to bacterial challenge independently of direct cell-cell signaling. Importantly, the ability of neuron-conditioned medium alone to preserve tight junction integrity identifies a previously unrecognized mechanism by which neurons contribute to BBB protection during infection.

**Figure 4:**
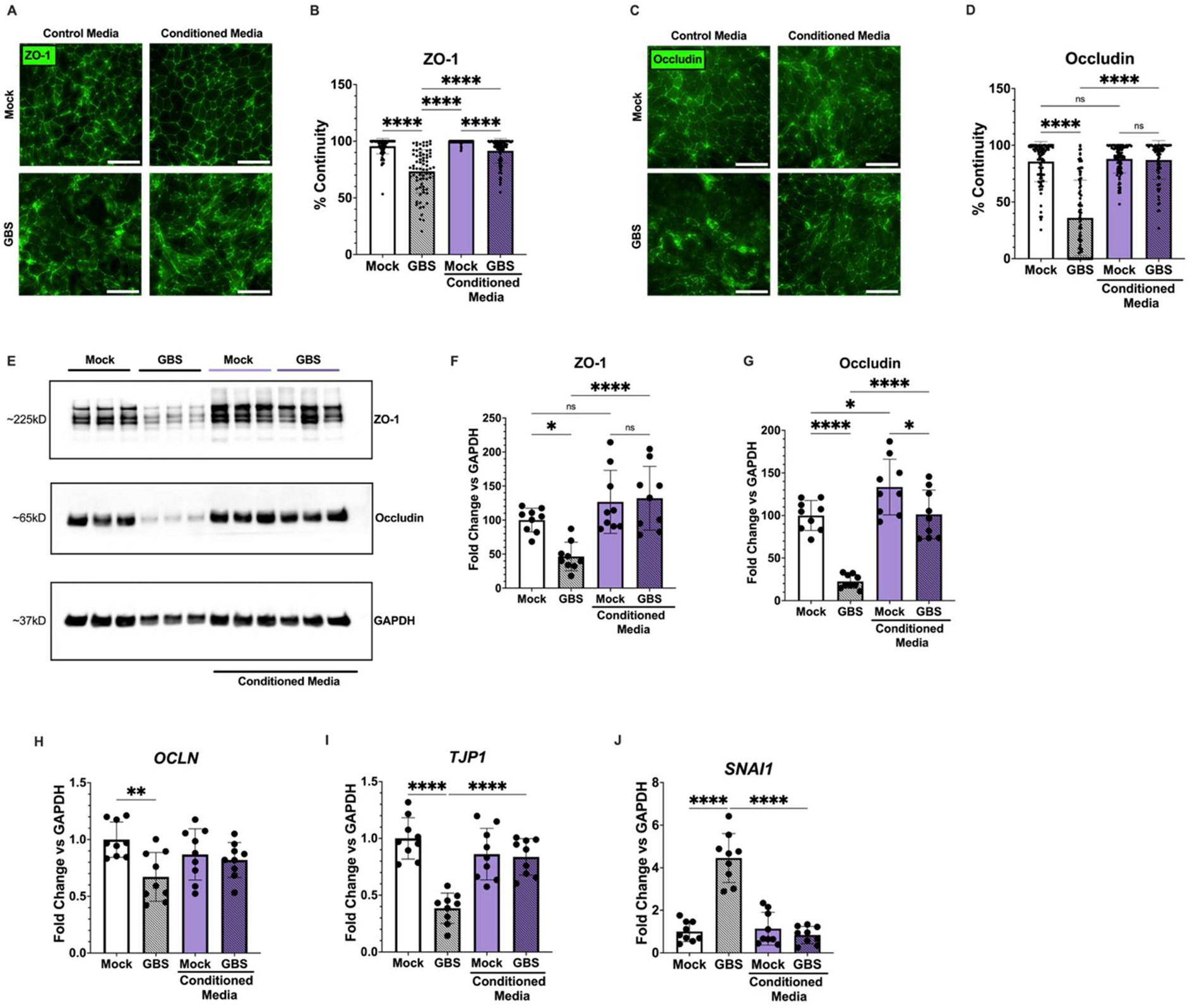
Neuron-conditioned medium preserves tight junction organization and barrier-protective responses during GBS infection. Representative immunofluorescence images of iBECs maintained in control or neuron-conditioned medium and subsequently infected with GBS. Cells were stained for the tight junction proteins (A) ZO-1 and (C) Occludin to assess junctional organization. Quantification of (B) ZO-1 and (D) Occludin junctional continuity using the Junction Analyzer Program (JAnaP). Treatment with neuron-conditioned medium increased tight junction continuity in GBS-infected iBECs compared with control-treated cells. (E) Representative immunoblot images of ZO-1 and Occludin protein expression in mock- and GBS-infected iBECs maintained in control or neuron-conditioned medium. (F-G) Densitometric quantification of ZO-1 and Occludin protein abundance normalized to GAPDH. Treatment with neuron-conditioned medium preserved tight junction protein expression during GBS infection. (H-J) Relative gene expression of (H) *OCLN*, (I) *TJP1*, and (J) *SNAI1* in mock- and GBS-infected iBECs maintained in control or neuron-conditioned medium. Neuron-conditioned medium mitigated infection-associated alterations in tight junction gene expression and prevented induction of *SNAI1*, a transcriptional repressor associated with tight junction disruption and BBB dysfunction. Together, these findings demonstrate that neuron-derived soluble factors are sufficient to preserve tight junction organization and promote barrier-protective molecular responses in iBECs during GBS infection. (B, D) Data are presented as percent continuity of individual cells from images acquired across three independent biological replicates in technical triplicate, (B) *n* = 73-105, (D) *n* = 84-98. (F-J) Data are presented as from three independent biological replicates in technical triplicate, (*n* = 9). Error bars represent mean ± SD. Scale bar represents 100 µm. Ordinary one-way ANOVA was used to determine significance; *p < 0.05, **p < 0.01, ****p < 0.0001, ns, non-significant.

## Discussion

To our knowledge, this study is the first to examine how neuron-endothelial communication influences BBB susceptibility to GBS infection using an isogenic human iPSC-derived neurovascular model. By combining iBECs and iNeurons derived from a common donor source, this model captures neurovascular interactions that are absent from traditional endothelial monoculture systems and enables investigation of how neuronal signaling shapes host-pathogen interactions at the BBB. Previous studies have established that neuronal co-culture promotes BBB maturation and enhances endothelial barrier properties, including improved tight junction organization and increased physiological relevance (23, 25, 47). Our findings extend these observations to a clinically relevant model of neonatal meningitis and suggest that neuron-endothelial communication remains functionally important during infection. These results indicate that BBB susceptibility to pathogen-mediated injury is influenced not only by endothelial-intrinsic responses but also by signals originating from neighboring neural cells.

Importantly, neuronal presence was associated with reduced endothelial susceptibility to GBS-induced dysfunction, supporting a previously underappreciated role for neurons in regulating BBB host defense. One interpretation of these findings is that neuronal signaling promotes endothelial barrier properties that are inherently less permissive to bacterial attachment and invasion. Alternatively, neuron-derived factors may directly inhibit endothelial signaling pathways involved in bacterial attachment, uptake, or maintenance of junctional integrity.

Human iPSC-derived BBB models provide important advantages for *in vitro* studies of bacterial meningitis compared with immortalized endothelial cell lines, which often lack robust BBB phenotypes and physiological responsiveness (1, 17, 48). Importantly, unlike many immortalized models, iBECs remain responsive to inductive signals from neighboring NVU cell types, including astrocytes and neurons derived from both primary and pluripotent stem cell sources, allowing investigation of multicellular interactions that regulate BBB function in health and disease (1, 23). These interactions have been shown to enhance BBB properties, including junctional organization and barrier integrity. However, how multicellular interactions influence host-pathogen dynamics during bacterial meningitis has yet to be explored. Our findings address this gap by incorporating neuronal components into the model and demonstrating their impact on BBB responses during GBS infection. These observations highlight the importance of incorporating physiologically relevant BBB models for elucidating mechanisms contributing to pathogen-mediated barrier disruption and for identifying potential therapeutic strategies aimed at limiting BBB disruption during meningitis. Collectively, our findings suggest that BBB protection during GBS infection is not solely an endothelial intrinsic process but instead reflects coordinated signaling within the NVU. This is consistent with emerging models in which NVU communication contributes to BBB resilience during neuroinflammation and pathological states (49). The observation that neuron-conditioned medium recapitulated many of the effects of co-culture suggests that soluble neuron-derived factors may be sufficient to promote endothelial resistance to bacterial injury. Further studies will be required to identify these factors and define the mechanisms by which neuronal signaling modulates endothelial responses to GBS.

Neurons are known to secrete a diverse array of soluble mediators including molecular chaperones, metabolic enzymes, and proteasome subunits, which have been implicated in processes such as neurogenesis, neurovascular communication, endothelial stability, and immune signaling (50–53). These observations support the hypothesis that neuron-derived factors contribute to the regulation of BBB integrity and endothelial responses during infection. The ability of neuron-conditioned medium to reproduce the protective effects observed in co-culture suggests that soluble neuronal signals are sufficient to enhance BBB resilience to GBS challenge. Future studies aimed at characterizing the neuronal secretome through proteomic and other unbiased discovery approaches will be essential for identifying the specific mediators responsible for this protective phenotype (54, 55). Defining these factors will provide important mechanistic insight into how neuronal signaling regulates endothelial function and may reveal novel therapeutic targets for preserving BBB integrity during bacterial meningitis. Limitations of this study should be considered for future investigation. Although the isogenic co-culture model captures important aspects of neuron-endothelial communication, it does not fully recapitulate the complexity of the *in vivo* neurovascular unit, which includes astrocytes, pericytes, microglia, and infiltrating immune cells, and dynamic vascular flow. Static culture conditions used here do not model the shear stress and biological cues experienced by brain endothelial cells *in vivo*, which may influence barrier function and host-pathogen interactions. In addition, while neuron-conditioned medium experiments support a role for soluble signaling, the identity of the responsible factors remain unknown. Furthermore, our study focused on a most clinically relevant hypervirulent GBS strain, COH1 (Serotype III, ST-17), and it remains to be determined whether similar neuroprotective mechanisms operate across diverse GBS lineages or other meningitis-causing pathogens. Addressing these limitations through more complex neurovascular models will be important for defining the broader relevance of neuronal regulation of BBB susceptibility during infection and for establishing whether neurovascular host defense mechanisms are conserved across bacterial meningitis pathogens.

In conclusion, this study introduces an isogenic, human-relevant BBB model as a platform for interrogating host-pathogen interactions during bacterial meningitis. Our findings demonstrate that neuronal-endothelial communication significantly influences the outcome of GBS infection at the BBB by reducing bacterial adherence and invasion, preserving tight junction organization, and mitigating barrier dysfunction. More broadly, these findings identify neuron-endothelial crosstalk as a previously unrecognized component of host defense at the BBB. Understanding how neuronal signaling enhances endothelial resistance to infection may reveal novel strategies for preserving barrier function and limiting bacterial invasion of the CNS. Identifying the neuron-derived soluble factors responsible for these protective effects will be critical for defining the molecular mechanisms underlying BBB resilience during infection and may reveal novel targets for adjunctive therapies in neonatal meningitis. Furthermore, patient-specific iPSC-derived neurovascular models provide a scalable and flexible platform for investigating how genetic background and disease state influence BBB function and susceptibility to infection. Such approaches may also be extended to other CNS infections and neurological disorders in which BBB dysfunction contributes to disease pathogenesis.

---

**Supplement Figure 1:**
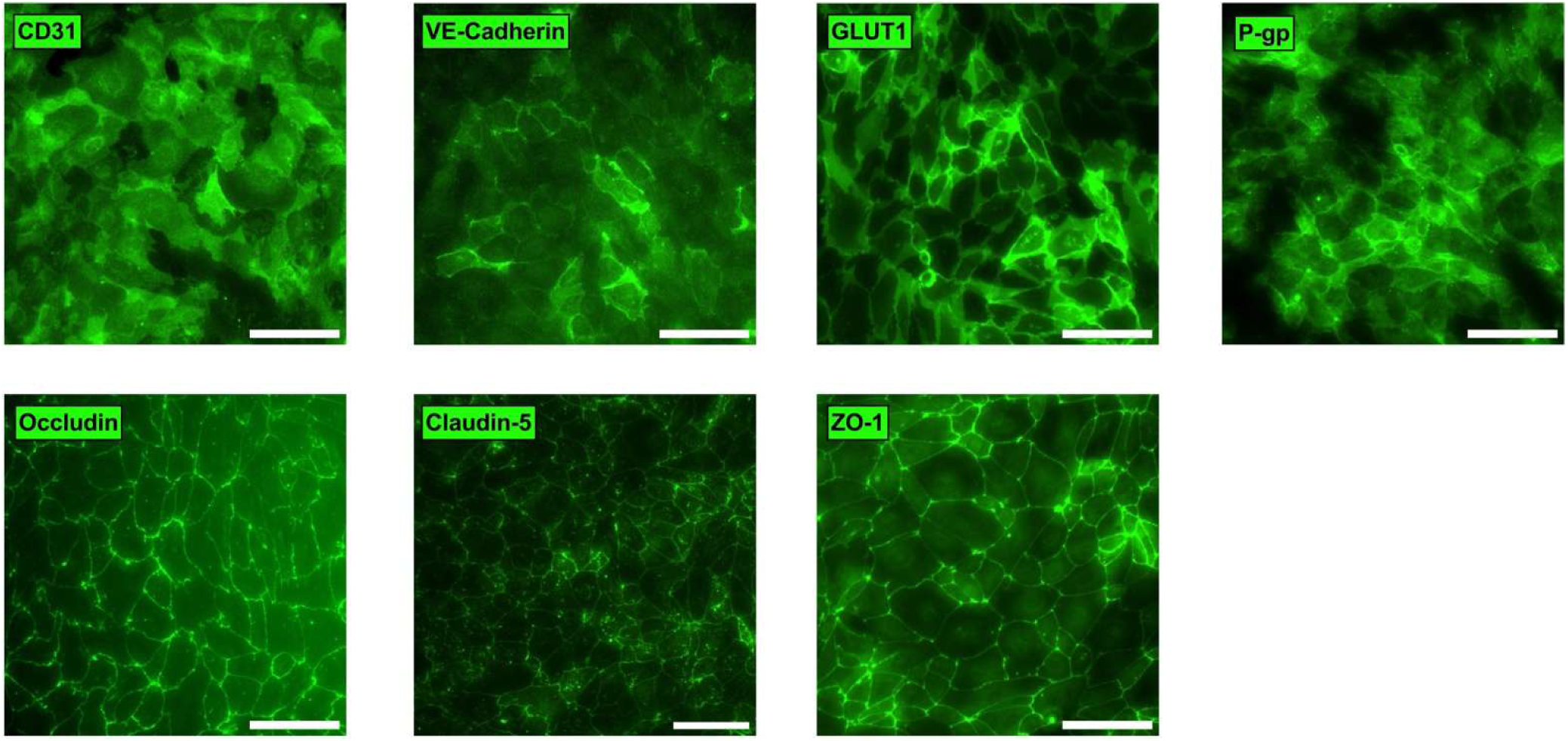
Characterization of iPSC-derived brain endothelial cells. Representative immunofluorescence images of iBECs stained for the endothelial and blood-brain barrier-associated markers CD31, VE-cadherin, GLUT1, P-gp, Occludin, Claudin-5, and ZO-1 following differentiation. Scale bar represents 100 µm.

**Supplement Figure 2:**
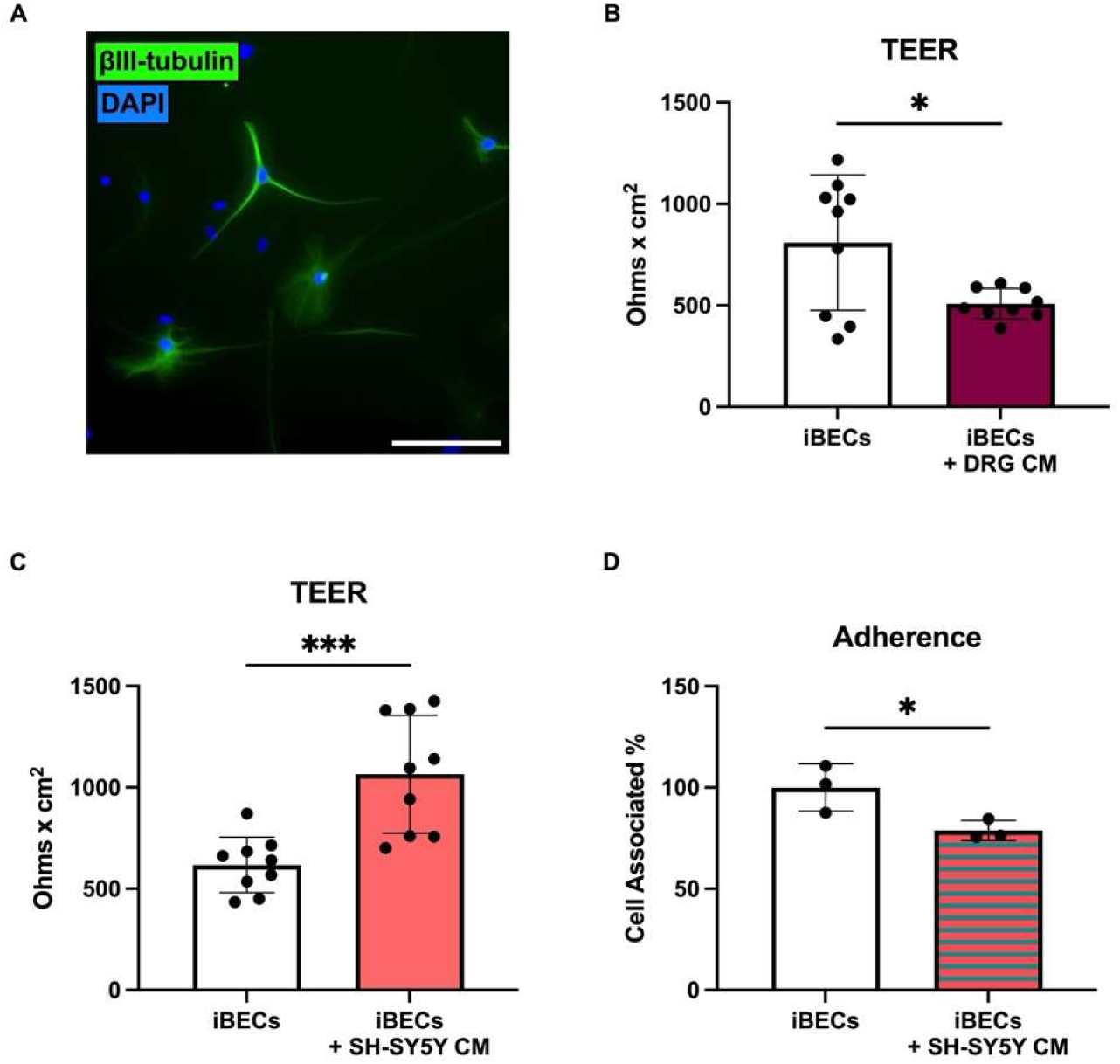
Effect of additional neuron models on iBEC barrier response. (A) Representative immunofluorescence image of primary mouse dorsal root ganglion (DRG) neurons stained for the neuronal marker βIII-tubulin, confirming neuronal identity. Scale bar represents 100 µm. (B-C) TEER measurements of iBEC monolayers following exposure to conditioned medium from (B) primary mouse DRG neurons or (C) SH-SY5Y neuroblastoma cells. Conditioned medium from SH-SY5Y cells significantly enhanced barrier integrity, whereas DRG neuron-conditioned medium significantly impaired barrier integrity, suggesting that the barrier-protective effects observed with EZ-Sphere-derived neurons may not be conserved across all neuronal populations. (D) Adherence of wild-type (WT) Group B *Streptococcus* (GBS) strain COH1 in iBECs following treatment with control or SH-SY5Y-conditioned medium shown in technical triplicate (N = 3). TEER data are presented as mean ± SD from three independent biological differentiations in technical triplicate, (*n* = 9). Statistical significance was determined by Student’s *t*-test; *p < 0.05, ***p < 0.001.

**Supplement Figure 3:**
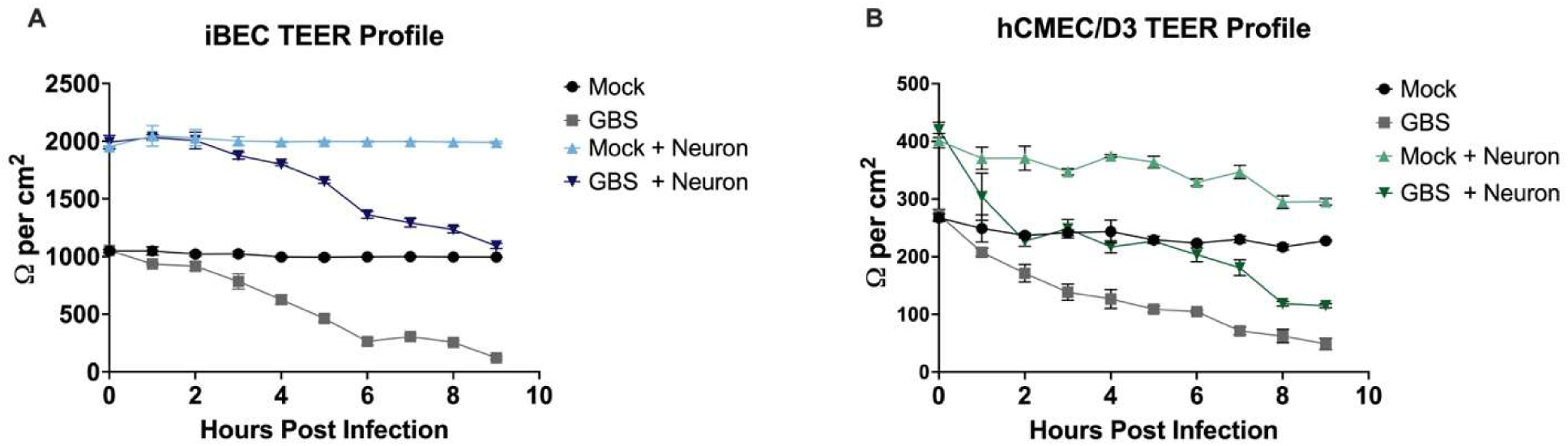
TEER measurements of endothelial monocultures and neuron co-cultures during GBS infection. TEER measurements of (A) iBECs and (B) hCMEC/D3s maintained as mock monocultures, GBS-infected monocultures, mock neuron co-cultures, or GBS-infected neuron co-cultures. TEER was monitored to assess changes in endothelial barrier integrity in response to neuronal co-culture and GBS infection. Data are presented in technical triplicate (N = 3).

**Supplement Figure 4:**
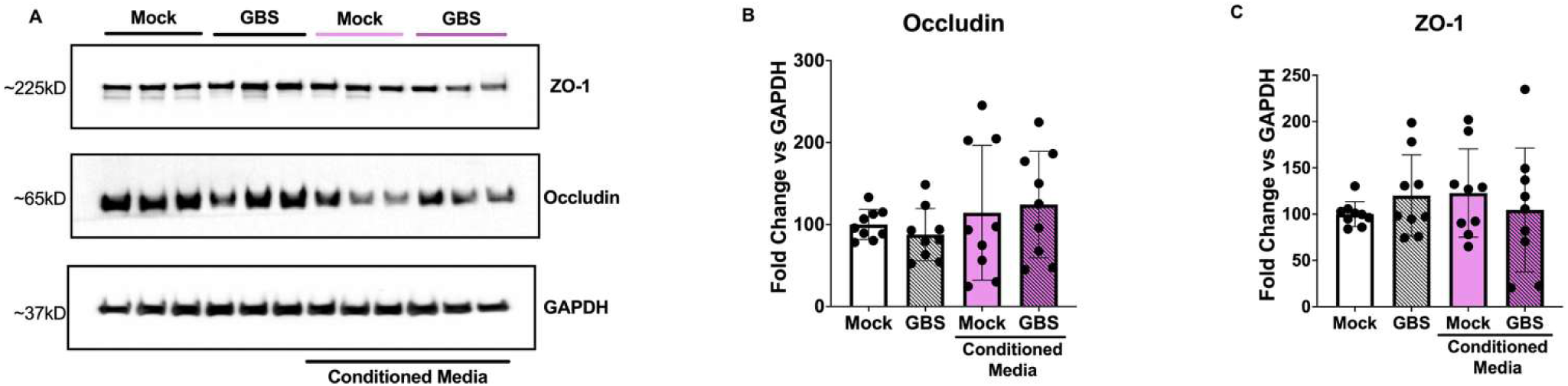
Neuron-conditioned medium does not significantly alter tight junction protein expression in hCMEC/D3 cells during GBS infection. (A) Representative immunoblot analysis of ZO-1 and Occludin protein expression in hCMEC/D3 cells maintained in control or neuron-conditioned medium under mock and GBS-infected conditions. Densitometric quantification of (B) Occludin and (C) ZO-1 protein abundance normalized to GAPDH. Neuron-conditioned medium did not significantly alter expression of either tight junction protein in GBS-infected hCMEC/D3 cells, indicating potential model-specific differences in endothelial responses to neuron-derived soluble factors. Data are presented as mean ± SD from three independent biological replicates performed in technical triplicate (*n* = 9). Statistical significance was determined by Ordinary one-way ANOVA.

## Acknowledgements

This research was funded by National Institute of Neurological Disorders and Stroke, grant number R15NS131921 awarded to B.J.K. M.D.B. received research support from The University of Texas System STARS program research support grant, and the Rita Allen Foundation Grant. The funders had no role in study design or decision to submit the work for publication. Diagrams were created in BioRender. The authors declare no conflicts of interest. The raw data supporting the conclusions of this article will be made available by the authors, without undue reservation.

